# *Drosophila* poised enhancers are generated during tissue patterning with the help of repression

**DOI:** 10.1101/052142

**Authors:** Nina Koenecke, Jeff Johnston, Qiye He, Julia Zeitlinger

**Author notes:** Present address: Department of Basic Medicine; School of Medical Sciences; Zhejiang University; Hangzhou, Zhejiang Province 310058, China (Qiye He). Corresponding author contact information: Julia Zeitlinger, 1000 East 50th St, Kansas City, MO 64110.

## Abstract

Histone modifications are frequently used as markers for enhancer states, but how to interpret enhancer states in the context of embryonic development is not clear. The poised enhancer signature, involving H3K4me1 and low levels of H3K27ac, has been reported to mark inactive enhancers that are poised for future activation. However, future activation is not always observed and alternative reasons for the widespread occurrence of this enhancer signature have not been investigated. By analyzing enhancers during dorsal-ventral (DV) axis formation in the *Drosophila* embryo, we find that the poised enhancer signature is specifically generated during patterning in the tissue where the enhancers are not induced, including at enhancers that are known to be repressed by a transcriptional repressor. These results suggest that, rather than serving simply as an intermediate step before future activation, the poised enhancer state may mark enhancers for spatial activation during tissue patterning. We discuss the possibility that the poised enhancer state is more generally the result of repression by transcriptional repressors.

## Introduction

Understanding the mechanisms by which cis-regulatory elements, or enhancers, activate transcription has been intensively studied for the last three decades, yet our knowledge remains incomplete (Shlyueva et al. 2014). As shown by ChIP-seq experiments, transcription factors may bind to thousands of putative enhancer regions in the genome (Moorman et al. 2006; Li et al. 2008), yet a large fraction of them are likely inactive. For example, transcription factors may bind to enhancers that have been primed by pioneer transcription factors but are not yet active (Zaret and Carroll 2011; Spitz and Furlong 2012) or they may be sequence-specific repressors that actively repress the enhancers to which they are bound (Sandmann et al. 2007; Zeitlinger et al. 2007). This raises the question of what types of enhancer states exist and how they help regulate the complex spatial and temporal expression patterns of genes during the development of multicellular organisms.

Good markers for enhancer states are the histone modifications found at the nucleosomes flanking enhancer regions. Most open enhancer regions are marked by histone mono-methylation on lysine 4 of histone H3 (H3K4me1), but only active enhancers carry lysine 27 acetylation on histone 3 (H3K27ac) (Creyghton et al. 2010; Ernst et al. 2011; Rada-Iglesias et al. 2011; Zentner et al. 2011; Bonn et al. 2012). Since some inactive enhancers show activation during later development, the combination of H3K4me1 along with low H3K27ac at inactive enhancers was termed the poised enhancer signature (Creyghton et al. 2010; Rada-Iglesias et al. 2011).

The mechanisms by which poised enhancers remain inactive and by which they become active under some conditions are poorly understood. For example, some studies have implicated the Polycomb-repressive mark H3K27me3 as a marker for poised enhancers (Rada-Iglesias et al. 2011), while others have not (Creyghton et al. 2010; Bonn et al. 2012). It is also possible that other mechanisms of repression might make enhancers susceptible to de-repression, thereby poising them for activation.

Poised enhancers are very common during the development of *Drosophila* and mammalian lineages, but their role in tissue patterning and lineage specification remains unclear. While originally described as being poised for future activation, this model is likely an oversimplification. The majority of enhancers become active without going through a poised state during prior developmental stages (Bonn et al. 2012; Choukrallah et al. 2015). Only a small fraction of poised enhancers are usually activated during lineage development (Rada-Iglesias et al. 2011; Bonn et al. 2012; Wamstad et al. 2012). Instead, many enhancers that are poised in a cell type are active in related cell types (Bonn et al. 2012; Junion et al. 2012; Wang et al. 2015). This not only questions the strict temporal model in which the poised enhancer state precedes enhancer activation, but also suggests a role for poised enhancers in tissue patterning.

A widespread role for poised enhancers in tissue patterning is consistent with large-scale DNase hypersensitivity (DHS) assays across a variety of cell types representing stages of human development (Stergachis et al. 2013). These data also show that enhancers are frequently accessible across broadly related cell types and only become active in specific lineages, raising the possibility that poised enhancers in embryonic tissues are predisposed for activation spatially, and that enhancer activation is regulated by signals that control pattern formation.

During tissue patterning, developmental signals (or morphogens) are often generated at and propagated from precise locations within the embryo, typically leading to the graded activation of signal transduction pathways and transcription factors across fields of cells (Briscoe and Small 2015). Depending on the strength of signaling, different target genes are activated, giving rise to distinct cell fates across the gradient. Activation of already accessible enhancers is a logical mechanism by which signal transduction pathways could mediate precise cellular responses to morphogens. The broad distribution of poised enhancers may ensure that a sufficient number of cells can respond to specific developmental signals in the appropriate manner, thus facilitating pattern formation.

While a function of poised enhancers in pattern formation is plausible, in many systems the hypothesis is difficult to test due to the scarcity and heterogeneity of embryonic tissues. To analyze a possible role for poised enhancers during pattern formation in the embryo, we used the tractable *Drosophila* dorso-ventral (DV) patterning as model system. In the *Drosophila* embryo, DV patterning begins with localized activation of the Toll (Tl) receptor by maternal components, which leads to the formation of a Dorsal (Dl) morphogen gradient and gives rise to at least three cell fates with distinct gene expression programs along the DV axis: mesoderm on the ventral side, neurectoderm in the lateral regions and dorsal ectoderm on the dorsal side (Hong et al. 2008) (Fig. 1a). For simplicity, we focused on the cell fates at the ends of the gradient, mesoderm and dorsal ectoderm.

**Figure 1.**
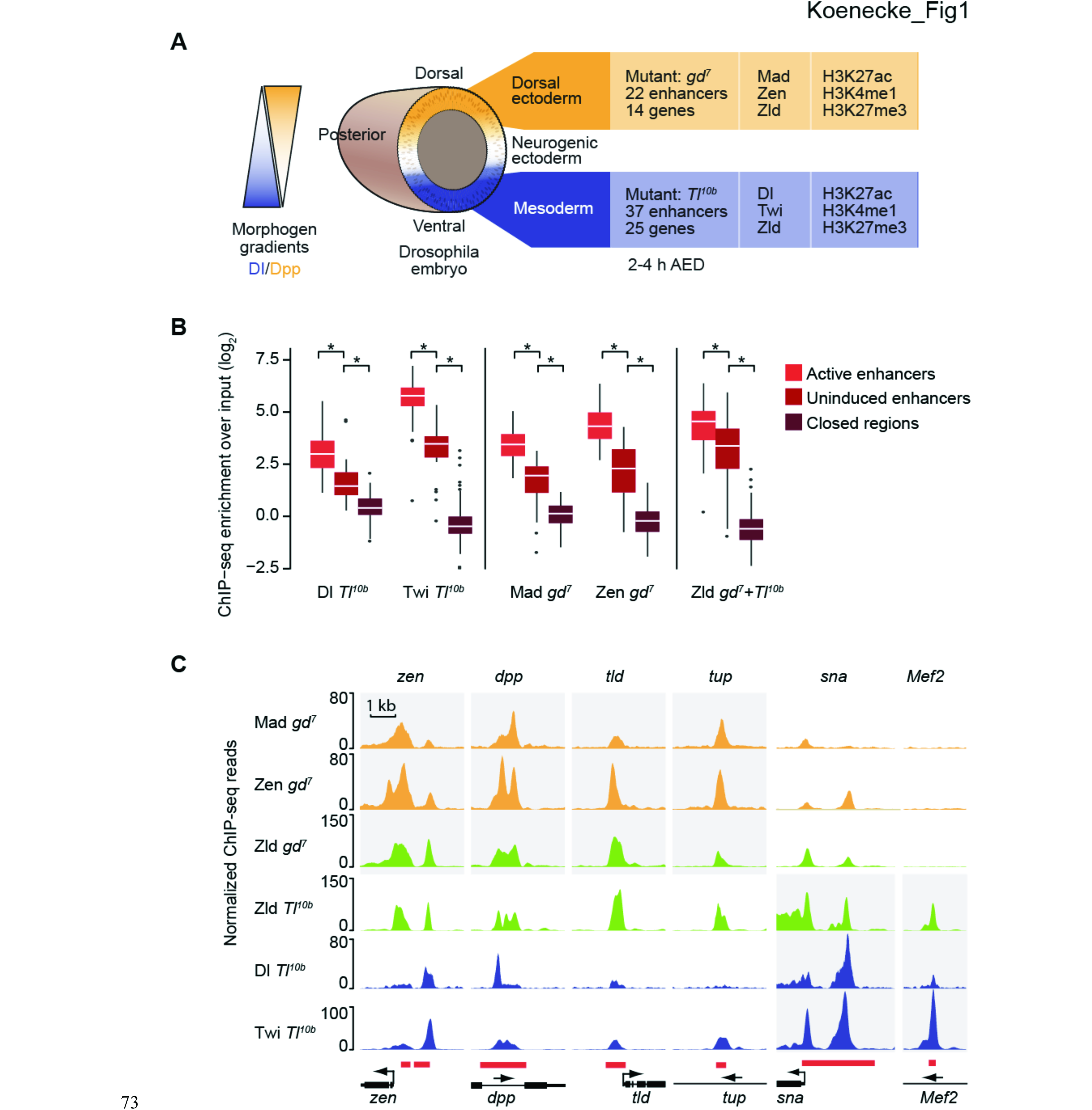
Intermediate levels of DV transcription factors at uninduced enhancers. **(A)** Overview of the model system of the dorsal-ventral (DV) patterning in the *Drosophila* embryo, in which homogenous cell fates can be obtained through mutants such as *gd^7^* and *Tl^10b^* and for which a large number of tissue-specific enhancers and their target genes are known. The ChIP-seq experiments performed in this study are also summarized. AED = after egg deposition **(B)** Boxplots of ChIP-seq data over input for the DV transcription factors Dl, Twi, Mad, Zen and Zld at known DV target enhancers. The ChIP-seq experiments were performed in either *gd^7^* or *Tl*^*10b*^, or both, dependent on where the transcription factor is expressed. Note that DV transcription factors occupy uninduced enhancers less than active enhancers but significantly more than closed regions, indicating that uninduced enhancers are accessible. Closed regions are 100 presumptive late enhancers that are inaccessible by DHS at early stages (Thomas et al. 2011) and are enriched for H3K27ac at later stages (see Methods for details). Active enhancers are MEs in *Tl^10b^* embryos or DEEs in *gd^7^* embryos. Uninduced enhancers are MEs in *gd^7^* embryos or DEEs in *Tl^10b^* embryos. Whiskers show 1.5 times the interquartile range and outliers are shown as dots. Asterisk indicates p < 10^−4^, using the Wilcoxon rank sum test. **(C)** ChIP-seq binding profiles of the transcription factors at four DEEs and two MEs (red boxes with target genes shown in black) illustrate higher binding at active enhancers (grey shading) but some degree of binding at uninduced enhancers.

The advantage of the *Drosophila* DV system is that large amounts of cells can be obtained from these two tissues without the need for cell sorting or tissue dissection. This is made possible by the availability of maternal mutants where all embryos in the progeny consist entirely of either mesodermal or dorsal ectodermal precursor cells (Schneider et al. 1991). In *Tl^10b^* mutant embryos, Dl activity is uniformly high (but not above wild-type levels) leading to mesodermal precursor fate. In *gd^7^* mutant embryos, Dl is not activated, resulting in uniformly high signaling activity of the fly BMP2/4 ortholog Decapentaplegic (Dpp) (but below wild-type maximum levels, see Ashe and Levine 1999) and the specification of dorsal ectodermal fate in the entire embryo. These mutants have frequently been used in the past because they allow the analysis of patterning across the Dl activity gradient (e.g. Stathopoulos et al. 2002; Zeitlinger et al. 2007; Holmqvist et al. 2012), and have helped DV patterning become one of the best-studied gene regulatory networks in development.

The DV patterning system also illustrates another important principle of pattern formation, the widespread use and requirement of sequence-specific transcriptional repressors. The extensive genetic screens in *Drosophila* have shown that transcriptional repressors are crucial for the correct interpretation of morphogen gradients, including DV patterning (Ip and Hemavathy 1997; Bier and De Robertis 2015; Briscoe and Small 2015). During DV patterning, Dl is able to specify three distinct cell fates because, in addition to its role as a transcriptional activator, it can also act as a repressor when certain additional repressive sequences in an enhancer are present next to a Dl motif (Pan and Courey 1992; Jiang et al. 1993). When Dl is converted into a repressor, it recruits co-repressors and histone deacetylases (Dubnicoff et al. 1997; Valentine et al. 1998; Chen et al. 1999; Flores-Saaib et al. 2001; Ratnaparkhi et al. 2006) and dominantly suppresses enhancer activation (Gray and Levine 1996; Dubnicoff et al. 1997). Three cis-regulatory sequences, those regulating *dpp, zerknüllt* (*zen*), and *tolloid* (*tld*), have been shown to be ventrally repressed by Dl, allowing spatially-restricted activation of these genes on the dorsal side of the embryo (Irish and Gelbart 1987; Rushlow et al. 1987; Ip et al. 1991; Huang et al. 1993; Kirov et al. 1994; Ratnaparkhi et al. 2006).

Using the DV patterning system, we analyzed the state of enhancers during patterning across tissues. We show that DV enhancers are indeed in a poised state in the tissue where they are not induced, including at the three loci that are known to be repressed. These enhancers are accessible to transcription factors, albeit at lower levels than in their active state, and marked by significant levels of H3K4me1 but low levels of H3K27ac. Their H3K27me3 levels were more variable. We find no evidence that these poised enhancers mark future activation. The poised enhancer signature is not present before DV patterning and DV enhancers are not open beyond DV patterning. This shows that the poised enhancer signature can specifically arise during patterning, in tissues where the enhancer is not induced. We discuss the possibility that the poised enhancer signature is a result of repression, and propose a model in which enhancer-bound repressors are a critical component of enhancer regulation with important mechanistic implications.

## Results

### Uninduced DV enhancers are accessible to transcription factors albeit at lower levels

To characterize the enhancer states during DV patterning, we first assembled a list of known DV enhancers that have been verified by transgenic *lacZ* reporter assays (see Supplemental Table S1). We identified 37 enhancers that exhibit activity in the mesoderm but remain uninduced in the dorsal ectoderm, and 22 enhancers that drive expression in the dorsal ectoderm but remain uninduced in the mesoderm (Fig. 1A, see Supplemental Material for a complete list and references). To validate our experimental system, we performed mRNA-seq experiments on *Tl^10b^* and *gd^7^* embryos at 2-4 h after egg deposition (AED), the time window during which DV patterning occurs (Stathopoulos et al. 2002; Zeitlinger et al. 2007). As expected, most genes that were assigned to a known DV enhancer were more highly expressed in the tissue in which the enhancer is active (Supplemental Fig. S1).

**Supplemental Figure 1.**
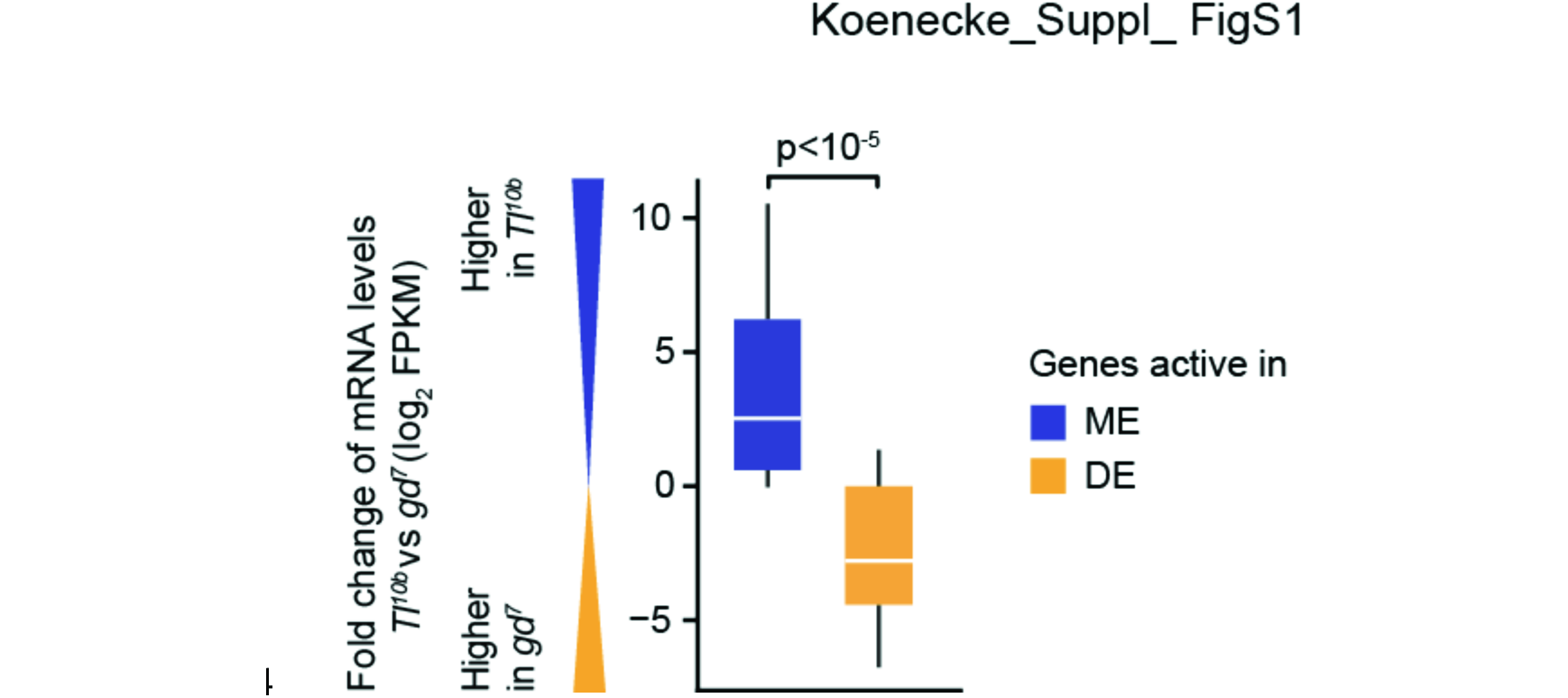
Analysis of transcript levels at DV enhancers across tissues. Boxplots of the fold change in transcript levels of DV genes between *Tl^10b^* embryos and *gd^7^* embryos (log_2_ FPKM). Whiskers show 1.5 times the interquartile range. Only confirmed target genes for the known DV enhancers were included (see Supplemental Material). Mesoderm enhancers (MEs), dorsal ectodermal enhancers (DEEs).

To measure the binding of transcription factors to DV enhancers in the active versus uninduced state during DV patterning, we performed ChIP-seq experiments in *Tl^10b^* and *gd^7^* embryos at 2-4 h AED. Replicate experiments were highly correlated (see Supplemental Material). We specifically analyzed DV transcription factors that are required for the cell fate specification of mesoderm and dorsal ectoderm (Fig. 1A). High Dl activity on the ventral side of the embryo induces Twist (Twi), which together with Dl activates mesodermal target genes (Jiang et al. 1991; Ip et al. 1992). We therefore analyzed Dl and Twi occupancy in *Tl^10b^* embryos and calculated their enrichments at active mesoderm enhancers (MEs), as well as at dorsal ectoderm enhancers (DEEs), which are actively repressed or remain uninduced (Fig. 1B left). As a control, we used a set of 100 presumptive late enhancers that are inaccessible (“closed”) at 2-4 h AED but are accessible and marked by H3K27ac in the late embryo (see Methods). Active enhancers had the highest levels of Dl and Twi, the “closed” control set had the lowest levels, and uninduced enhancers had statistically distinct intermediate levels (Fig. 1B left).

Dorsal ectodermal fate is induced by Dpp signaling, which activates the transcription factors Mothers against dpp (Mad) and Zen (Rusch and Levine 1997; Lin et al. 2006). We therefore analyzed the occupancy of Mad and Zen in *gd^7^* embryos and found that their occupancy at both active DEEs and uninduced MEs was also significantly higher than at the “closed” control enhancers (Fig. 1B, middle). Again, their occupancy at uninduced enhancers was significantly lower than at active enhancers (Fig. 1B, middle), further supporting the hypothesis that uninduced enhancers are bound by transcription factors, but to a lesser extent than active enhancers.

The observation that uninduced DV enhancers are bound by transcription factors suggests that these enhancers have been primed by a pioneer transcription factor. A potential pioneer transcription factor is Zelda (Zld, encoded by the *zld* gene also known as *vfl*), which is present ubiquitously in the *Drosophila* early embryo and primes enhancers even before DV patterning begins (Liang et al. 2008; Harrison et al. 2011; Nien et al. 2011). While Zld is required to make some DV enhancers accessible (Yanez-Cuna et al. 2012; Foo et al. 2014; Schulz et al. 2015; Sun et al. 2015), it is not known whether Zld remains bound to uninduced enhancers at the same level as at active enhancers.

We therefore analyzed the occupancy of Zld at active, uninduced and closed enhancers. Since Zld is present in both tissues, we merged the results for all active and all uninduced enhancers from both tissues (Fig. 1B right). We found that uninduced enhancers remain highly bound by Zld albeit at slightly lower levels than at active enhancers. The closed regions that we used as controls were not bound by Zld or bound at very low levels. This suggests that Zld specifically primes early enhancers and that it primes them in the entire embryo, whether or not the enhancers are induced.

Examples of transcription factor binding patterns at DV enhancers are illustrated in Fig. 1C. The three DEEs *zen, dpp* and *tld* show high occupancy of Zld, Mad and Zen in *gd^7^* embryos where they are active, as expected. In *Tl^10b^* mutants, where these enhancers are repressed, they are occupied by Zld, Dl and Twi. This is consistent with Zld’s role as pioneer factor and Dl’s role as repressor at these enhancers. However, these and other DEEs such as *tup* are also occupied by Twi to some degree, although Twi is an activator and has no known role in regulating these enhancers. This suggests that the DEEs are to some degree accessible to transcription factors in the tissue in which they are not induced, presumably due to the pioneering activity of Zld.

A similar pattern was observed for MEs. In *Tl^10b^* mutants, the *sna* enhancer is highly occupied by Dl and Twi, which are required for activation (Ip et al. 1992). However, even in the dorsal ectodermal tissue of *gd^7^* mutants, in which *sna* is not expressed, the enhancer is occupied by Zld, as well as Mad and Zen. Consistent with Zld being critical for enhancer access, in the rare case where Zld does not occupy an uninduced enhancer, other transcription factors are also not bound (see *Mef2* in Fig. 1C).

Taken together, these results suggest that uninduced enhancers are frequently primed and bound by transcription factors, albeit to a lower degree than in the active state. This level of accessibility might allow these enhancers to be inactive but responsive to changes in signaling and transcription factor activity.

### Uninduced enhancers are marked by H3K4me1 and low H3K27ac and thus carry a poised enhancer signature

Having identified three distinct enhancer states, we next investigated their histone modification status. We performed ChIP-seq experiments with antibodies against H3K27ac and H3K4me1 in both mutant embryos and calculated the enrichment (± 500 bp from enhancer center) at active and uninduced enhancers (again each from both mutants), as well as “closed” enhancers as a control.

Uninduced enhancers had overall significantly higher levels of H3K27ac as compared to closed control regions (Fig. 2A, p < 10^−6^, Wilcoxon rank sum test) but their levels were significantly lower than at active enhancers (Fig. 2A, p < 10^−6^, Wilcoxon rank sum test). Indeed, when we plotted the relative difference for each enhancer between the two tissues, the difference in H3K27ac levels between active and uninduced enhancers became more significant (p < 10^−9^) (Fig. 2B). This suggests that uninduced enhancers have low levels of H3K27ac, and that the levels significantly increase when the enhancers are active.

**Figure 2.**
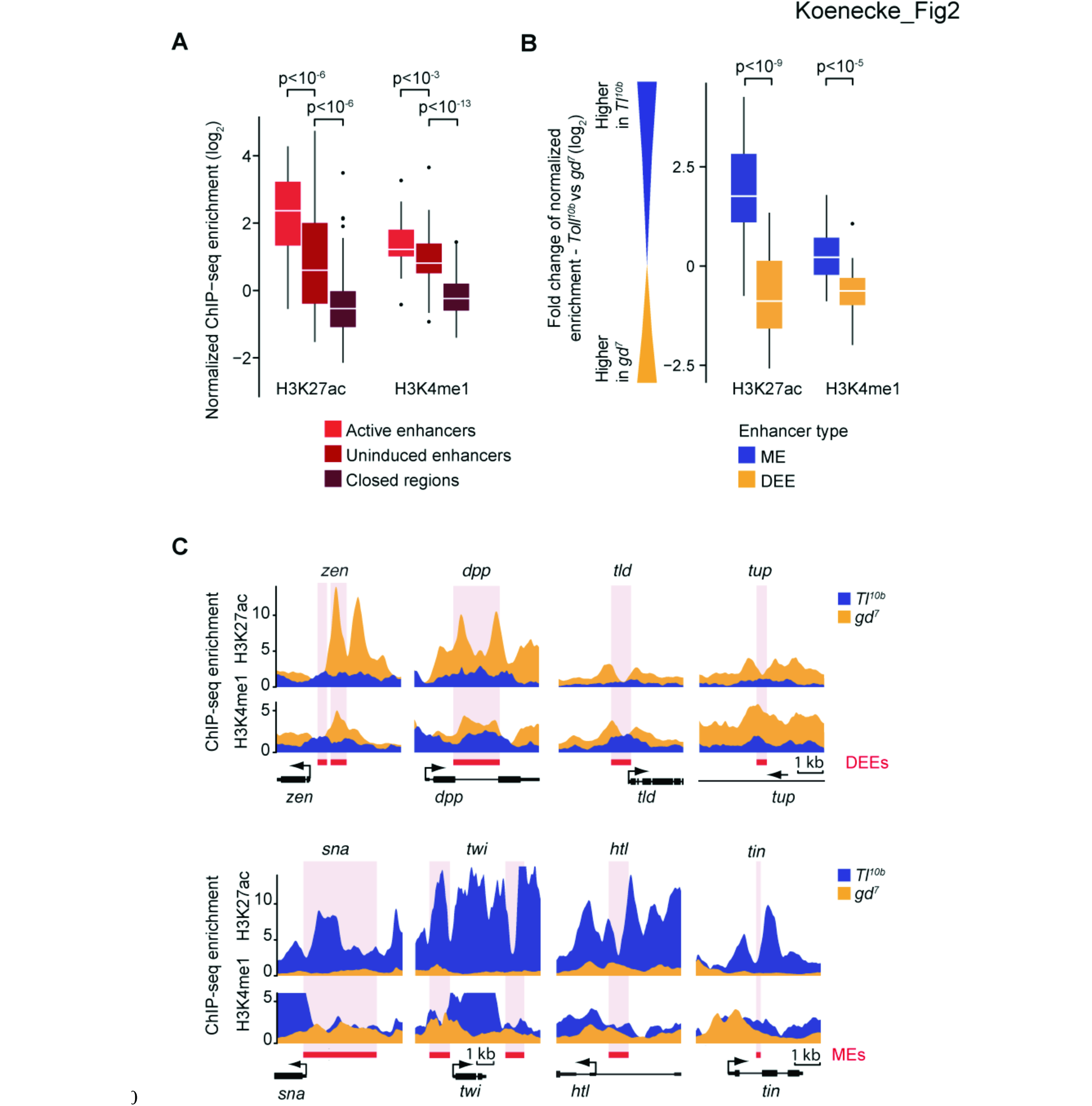
Figure 2. The histone modifications at uninduced enhancers resemble the poised enhancer signature. **(A)** Boxplots of normalized H3K27ac and H3K4me1 ChIP-seq enrichments over input show that all uninduced DV enhancers (n=59, when summed for both mutants) have lower H3K27ac enrichment levels than the same enhancers in the active state, yet the levels of H3K4me1 are significantly above closed regions (n=100, same as in Fig. 1), consistent with a poised enhancer signature. Whiskers show 1.5 times the interquartile range and outliers are shown as dots. Significance between enhancer groups was determined using the Wilcoxon rank sum test. **(B)** Boxplots of the fold-change of normalized histone modification ChIP-seq enrichments between mutant embryos show that H3K27ac and H3K4me1 levels are higher at active enhancers versus uninduced enhancers: the majority of mesodermal enhancers (MEs) (blue) have higher H3K27ac enrichment in the *Tl^10b^* mutant than in the *gd^7^* mutant (thus log_2_ *Tl^10b^* - log_2_ *gd^7^* above 0), while the inverse is true for dorsal ectodermal enhancers (DEEs) (yellow). Significance between enhancer types was determined using the Wilcoxon rank sum test. **(C)** Binding profiles of histone modification ChIP-seq enrichments show the higher enrichment of H3K27ac and H3K4me1 when the enhancer is active. Thus, at the four dorsal ectoderm enhancers (DEEs), the levels are higher in *gd7* (yellow), while at four mesoderm enhancers (MEs), the levels are higher in *Tl^10b^* (blue). The red box and the pink stripe show the position of the enhancers and the black arrow indicates the position and orientation of transcription start sites.

When we analyzed H3K4me1 levels, we found that uninduced enhancers also have H3K4me1 significantly above the levels of the control (Fig. 2A, p < 10^−13^, Wilcoxon rank sum test), consistent with a poised enhancer signature. However, H3K4me1 enrichments were slightly lower in the uninduced state than in the active state (Fig. 2A, p < 10^−3^, Wilcoxon rank sum test). This small but consistent difference became more significant when analyzing the relative difference in H3K4me1 at enhancers (p < 10^−5^) (Fig. 2B). Furthermore, close examination of the profiles of H3K4me1 at individual enhancers confirms this trend (see Fig. 2C). However, the difference is small relative to the difference between closed and uninduced regions, consistent with H3K4me1 being a marker for both poised and active enhancers.

Finally, we specifically examined whether the known enhancers repressed by Dl (*zen*, *dpp* and *tld* in Fig. 2C) had a characteristic histone modification signature distinct from other uninduced enhancers. The histone signature of H3K4me1 and low H3K27ac at repressed enhancers was indistinguishable (see other examples in Fig. 2C). Thus, the poised enhancer signature is also characteristic for enhancers regulated by transcriptional repressors. Whether there is a histone modification that is specifically associated with transcriptional repressors is not known. H3K27me3 is a well-studied repressive mark but it is deposited by Polycomb group proteins, which are not known to associate with sequence-specific transcriptional repressors (Simon and Kingston 2013).

### H3K27me3 is not a good marker for uninduced enhancers or sequence-specific repressors

Since the Polycomb-repressive mark H3K27me3 has been observed at poised enhancers (Rada-Iglesias et al. 2011) or repressed enhancers (Bonn et al. 2012), we tested whether H3K27me3 is found at DV enhancers and whether its presence correlates with a specific enhancer state. Polycomb group proteins typically regulate developmental genes (Schwartz et al. 2006; Tolhuis et al. 2006; Oktaba et al. 2008) but have not previously been implicated in embryonic DV patterning because their mutants are difficult to analyze in the early *Drosophila* embryo (Pelegri and Lehmann 1994).

When we analyzed H3K27me3 ChIP-seq data in *Tl^10b^* and *gd^7^* embryos, we found that H3K27me3 is present at DV enhancers, but at remarkably variable levels. Some enhancers had very high levels of H3K27me3, while more than half of them had no enrichment above background (Fig. 3A). Despite the variance, however, there was a significant trend for enhancers to have higher H3K27me3 levels in the uninduced versus active state (Fig. 3B, p < 10^−2^), consistent with previous findings (Bonn et al. 2012).

**Figure 3.**
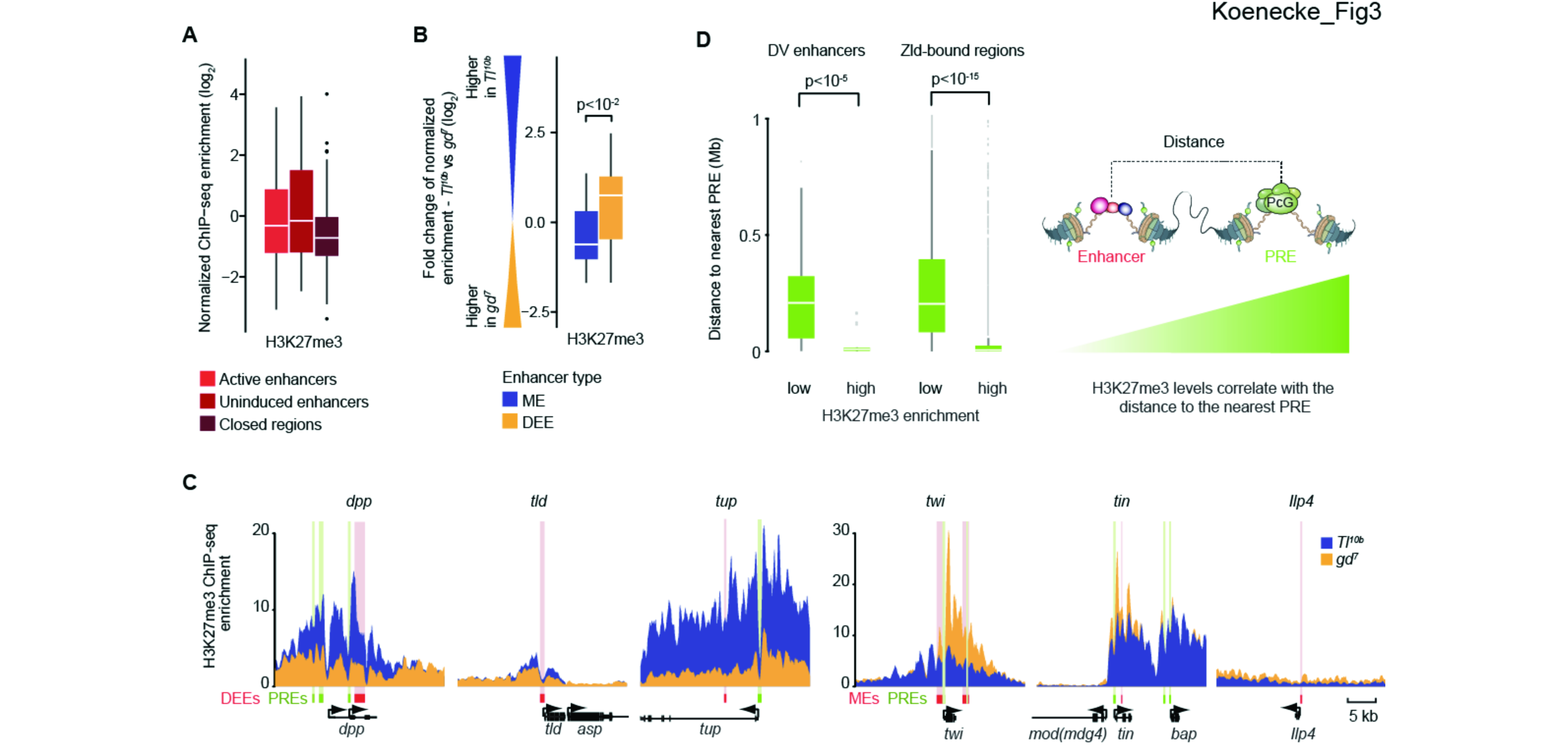
H3K27me3 levels are higher at uninduced enhancers but correlate more strongly with distance to the nearest PRE. **(A)** Boxplots of H3K27me3 show that a wide range of different levels are found at DV enhancers both in the active and uninduced state. **(B)** A relative plot of H3K27me3 levels between states shows that H3K27me3 levels at individual enhancers tend to be higher in the uninduced state versus active state. Significance was determined using the Wilcoxon rank sum test. **(C)** H3K27me3 ChIP-seq enrichment profiles for 3 dorsal ectoderm enhancers (DEEs) and 3 mesoderm enhancers (MEs) illustrate clear differences between mutants (yellow versus blue). H3K27me3 enrichment levels are highest near putative Polycomb response elements (PREs, green). Enhancers are shown as red box with pink shading. **(D)** Boxplots showing the distance of enhancers to the nearest PRE, dependent on whether they have low or high H3K27me3 enrichment levels. For DV enhancers with low H3K27me3 levels (n= 39), the distances are much larger than for those with high H3K27me3 levels (n=20). This is also true for Zld-bound regions (n=13,814), which includes a large number of putative early *Drosophila* enhancers. Putative PREs are defined as overlapping Pc and GAGA regions (see Supplemental Material for details). Whiskers show 1.5 times the interquartile range and outliers are shown as dots.

Examination of individual DV enhancers confirms clear differences in H3K27me3 levels between the uninduced and active state in regions where the levels of H3K27me3 are high (Fig. 3C). However, H3K27me3 marks are distributed over broad regions, as expected (Schwartz et al. 2006; Tolhuis et al. 2006); the differences in H3K27me3 include the transcribed regions and thus are not specific to DV enhancers (Fig. 3C). This questions whether an enhancer’s state directly regulates the surrounding levels of H3K27me3 or may instead affect H3K27me3 levels more indirectly through its effect on gene activation. Indeed, an anti-correlation between and H3K27me3 and transcriptional status has been observed previously (Klymenko and Muller 2004; Papp and Muller 2006; Tolhuis et al. 2006; Gaertner et al. 2012). This supports the traditional model in *Drosophila* in which gene activation by Trithorax group proteins counteracts Polycomb repression and reduces H3K27me3 (reviewed in Geisler and Paro 2015).

If gene activation reduces H3K27me3, what determines whether H3K27me3 is present in that region in the first place? Broad regions of H3K27me3 are catalyzed from specific nucleation sites in the DNA called Polycomb Responsive Elements (PREs) (Simon et al. 1993; Muller and Kassis 2006). In *Drosophila*, Polycomb group proteins are recruited to PREs by a combination of DNA-binding factors, including GAGA factor (*Trithorax-like* or *Trl*) (Strutt et al. 1997). We therefore identified high-confidence PREs through the co-occupancy of GAGA factor, which is not specific for PREs but gives high signal in ChIP experiments, and Polycomb (Pc) itself, which is indirectly bound to DNA but which is highly specific for PREs (Schuettengruber et al. 2009; Schuettengruber et al. 2014).

If the levels of H3K27me3 at enhancers depend on nearby PREs, we expect that DV enhancers with high H3K27me3 levels will be located closer to PREs than those without. Indeed, DV enhancers with H3K27me3 levels above 2-fold enrichment have PREs that are relatively close (median distance is less than 10 kb), while DV enhancers without H3K27me3 enrichment have PREs that are much further away (median distance is ~200 kb (Fig. 3D, p < 10^−5^, Wilcoxon rank sum test). The correlation between PREs and H3K27me3 can also be observed at individual DV enhancer regions, where the levels of H3K27me3 often peak close to PREs (Fig. 3C). Finally, the correlation between PREs and H3K27me3 is not specific for DV enhancers since the same trend was observed for all Zld-bound regions, which include most early enhancers (Fig. 3D). These results strongly support the traditional model that high levels of H3K27me3 depend on nearby PREs.

The anti-correlation between gene activation and H3K27me3 suggests that active enhancers can reduce the H3K27me3 levels deposited by nearby PREs. To consider alternative models, we also probed the possibility that repressors at enhancers might directly promote H3K27me3 deposition. However, the known Dl-repressed enhancers did not stand out in their H3K27me3 profile as compared to other uninduced enhancers (Fig. 3C). For example, the Dl-repressed *dpp* enhancer has very high levels of H3K27me3 in the repressed state, while another Dl-repressed enhancer, that of *tld*, has much lower levels. Furthermore, high levels are also observed at enhancers that are not repressed by Dl, including *tup*. Thus, while the levels of H3K27me3 correlate with the presence of PREs, they do not correlate with Dl-dependent repression. While we cannot rule out a subtle role for repressors in modulating H3K27me3 levels, our data suggest that the strongest determinants of H3K27me3 levels are nearby PREs and lack of gene activation. Therefore, H3K27me3 cannot be considered a specific marker for uninduced or repressed enhancers.

### Poised DV enhancers are specifically generated during tissue patterning and are not poised for future activation

Our results so far suggest that uninduced enhancers have a histone signature that is indistinguishable from the poised enhancer signature described in mammals, with or without H3K27me3. This raises the question whether the *Drosophila* DV enhancers are at some point poised for future activation.

We first considered the possibility that DV enhancers are poised prior to activation, when the enhancers are primed by Zld before Dl-dependent transcription begins. Based on a careful time-course analysis (Li et al. 2014), however, the primed DV enhancers do not show the poised signature since they gradually accumulate H3K27ac but are not yet marked with H3K4me1 or H3K27me3 before DV patterning takes place (Fig. 4A). This suggests that the DV enhancers do not have any poised enhancer signature when they are primed prior to activation.

**Figure 4.**
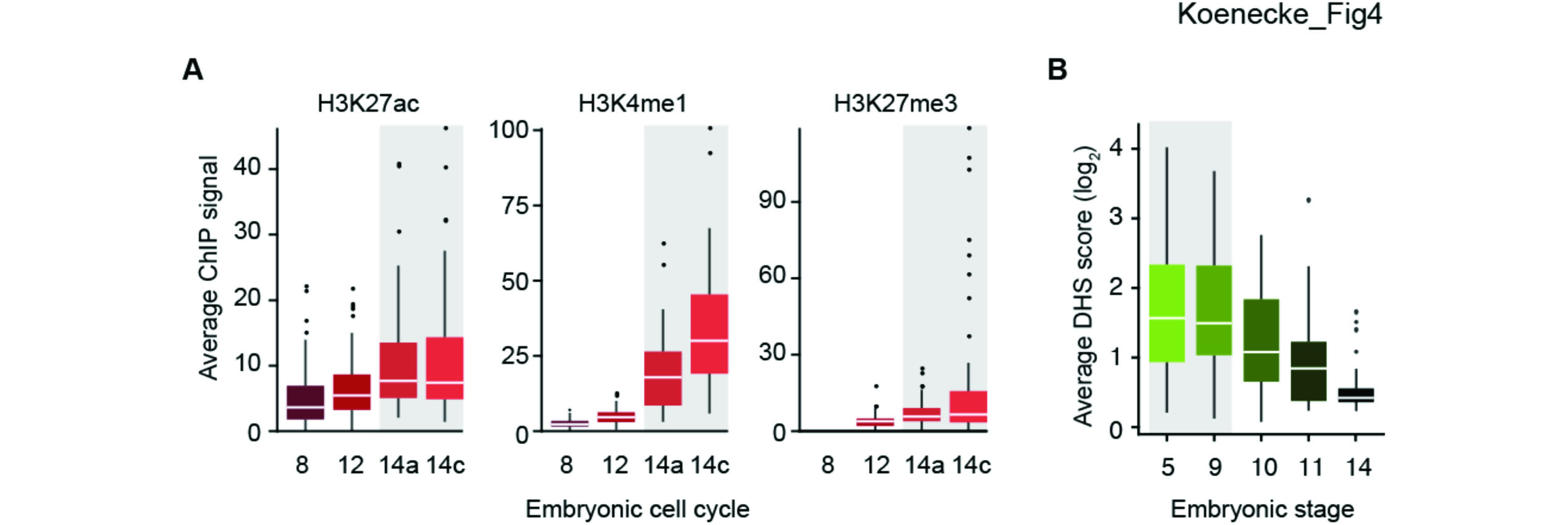
DV enhancers are not poised for future activation. (A) Histone modification levels at DV enhancers during the maternal-to-zygotic transition (Li et al. 2014) show that H3K27ac levels are accumulating early and gradually, thus some H3K27ac is present during enhancer priming by Zld. In contrast, H3K4me1 and H3K27me3, which mark poised enhancers, are only detectable after Dl-dependent transcription begins at stage 5 (cell cycle 14a and 14c, marked as grey box). **(B)** Boxplots of DNase I hypersensitivity (DHS) at DV enhancers during embryogenesis show that all DV enhancers are most accessible during stages 5 and 9 when DV patterning takes place (shaded in grey) and become less accessible at subsequent stages. The DHS score is the average signal per base derived from the data by Thomas et al. (2011). Whiskers show 1.5 the interquartile range and outliers are shown as dots.

We next considered whether the DV enhancers are poised for activation beyond DV patterning during later stages of embryogenesis. This seems unlikely since enhancers are in the vast majority stage-specific. To nevertheless test the possibility, we analyzed DNase I hypersensitivity (DHS) data across embryogenesis (Thomas et al. 2011). We found that DV enhancers are most accessible during DV patterning (stages 5 and 9), when they are active, and become less accessible at subsequent stages (Fig. 4B). This argues against additional roles of these enhancers past DV patterning.

Taken together, our analysis suggests that the poised enhancer signature is specifically generated during DV patterning at uninduced enhancers. There is no evidence that it precedes enhancer activation, arguing that it marks spatial rather than temporal regulation in our system.

## Discussion

### The poised enhancer signature as a marker for spatial enhancer regulation

We found that DV enhancers acquire the poised enhancer signature (low H3K27ac, some H3K4me1) specifically during tissue patterning (model in Fig. 5). Before DV patterning, these enhancers are primed by the pioneer transcription factor Zld and have a very different enhancer signature (some H3K27ac but no H3K4me1). It is unclear whether this enhancer signature is typical for primed enhancers since the priming occurs during the maternal-to-zygotic transition. Nevertheless, it clearly shows that the poised enhancer signature does not precede enhancer activation in the DV system and thus is specifically generated in the tissue in which the enhancers are not activated. During subsequent stages, the DV enhancers close again, perhaps because key transcription factors such as Zld are no longer present (Kanodia et al. 2012). It is also possible that repressive chromatin modifying complexes help to decommission enhancers to reduce their activity in subsequent developmental programs (Whyte et al. 2012).

**Figure 5.**
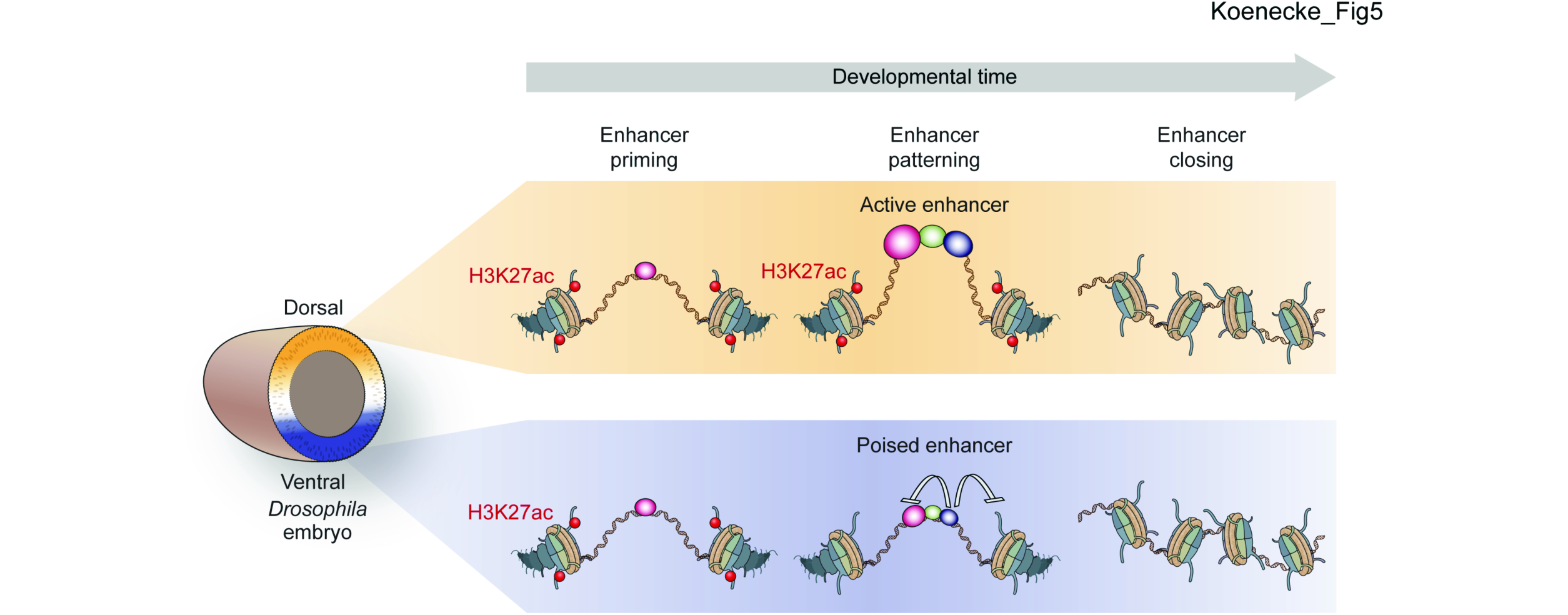
Summary model showing the poised enhancer signature specifically arising during tissue patterning in the Drosophila DV system. Before DV patterning begins in the *Drosophila* embryo, DV enhancers are primed by the pioneer transcription factor Zld and have low levels of H3K27ac. During DV patterning, DV enhancers may be active in one tissue but repressed by sequence-specific repressors in another tissue. These repressors recruit histone deacetylases, remove H3K27ac and produce the poised enhancer signature. After DV patterning is complete, DV enhancers gradually close, thus enhancers with the poised enhancer signature also close and are not poised for future activation.

This suggests that the poised enhancer signature should not be interpreted as “poised for future activation” but rather represents a “poised state”, one that would lead to activation in the presence of the right developmental signals. Since the “poised enhancer” is accessible to transcription factors, it can read out the activity of appropriate signal transduction pathways and respond to them. Therefore, enhancers may be in a poised state for some time during development to remain signal-responsive and allow cells to adjust to changes in signals from surrounding cells during pattern formation. However, in the absence of appropriate signals, a poised enhancer may not become active and instead may proceed directly to a closed state.

### A role for repressors in keeping poised enhancers inactive

We found that the three DV enhancers that are actively repressed by Dl have the poised enhancer signature. This raises the possibility that sequence-specific repressors actively help generate the poised enhancer signature and prevent these enhancers from becoming active.

In support of this hypothesis, the poised enhancer signature fits strikingly well with previous mechanistic studies on repression on individual loci in *Drosophila*. Transcriptional repressors such as Dl have been reported to reduce the occupancy of transcription factors and remove histone acetylation through the recruitment of co-repressors and histone deacetylases (Chen et al. 1999; Kulkarni and Arnosti 2005; Sekiya and Zaret 2007; Winkler et al. 2010; Li and Arnosti 2011). Thus, the low levels of H3K27ac and the reduced access to transcription factors that we observe for *zen, dpp* and *tld*, must be to some extent the result of Dl-mediated repression.

An even more intriguing hypothesis is that the poised enhancer signature is generally the product of enhancer-bound repressors. This would explain why the Dl-repressed enhancers did not stand out in their histone modification signature as compared to other uninduced enhancers. There are many sequence-specific repressors that modulate DV patterning, including Snail (Kosman et al. 1991; Leptin 1991), Capicua (Jimenez et al. 2000; Helman et al. 2012), Suppressor of Hairless (Morel and Schweisguth 2000; Ozdemir et al. 2014) and Schnurri (Crocker and Erives 2013). Thus, it is feasible that repressors play a central role in keeping enhancers inactive in the absence of activation. Below, we discuss a number of reasons why this is not only plausible but also an attractive model.

Based on ChIP-seq data, regions of open chromatin are surprisingly susceptible to unspecific transcription factor binding (Moorman et al. 2006; MacArthur et al. 2009). Many transcription factors have strong activation domains, putting accessible enhancer regions at risk for unwarranted activation. For example, Zld has high transactivation potential and can recruit the histone acetyl transferase CBP that mediates H3K27ac (Hamm et al. 2015; Stampfel et al. 2015), consistent with H3K27ac being present during enhancer priming by Zld (Li et al. 2014). Strikingly, we showed that Zld is still bound to DV enhancers during DV patterning, yet these enhancers have no or low H3K27ac and remain uninduced in parts of the embryo. The simplest explanation for this observation is that activation by Zld is repressed or “quenched” by repressors in these cells. Thus, repressors would serve to remove the histone acetylation that Zld induced during enhancer priming and prevent the accumulation of this activating mark throughout DV patterning.

Another reason is that the pattern by which poised enhancers occur during lineage development is consistent with the expected widespread use of repressors in signaling and tissue patterning. In addition to sequence-specific repressors employed during tissue patterning, most developmental signal transduction pathways have their own dedicated mechanism to repress target genes in the absence of signaling activity (Barolo and Posakony 2002; Affolter et al. 2008). The fact that these signal transduction pathways are highly conserved across evolution supports the notion that repression is an integral part of enhancer regulation.

### Mechanistic implications for poised enhancers with repressors

Finally, the involvement of repressors in keeping poised enhancers inactive has important mechanistic implications and predictions that have not been discussed to our knowledge. An active battle between activators and repressors in controlling histone acetylation at the poised state implies a monocycle between opposing enzymes, thus acetylation by acetyl transferases and deacetylation by deacetylases. Analogous to phosphorylation-dephosphorylation dynamics found at some enzymes, such monocycles can create switch-like behaviors and were therefore termed zero-order ultrasensitivity (Goldbeter and Koshland 1981; Ferrell and Ha 2014). In other words, repressors could make enhancers ultrasensitive in their response to activation signals.

Such zero-order ultrasensitivity predicts that a repressed enhancer can be very sensitive to activation, so that only a small amount of activation signal can lead to significant induction (Melen et al. 2005; Ferrell and Ha 2014). This is particularly important in the response to morphogen gradients, where a certain threshold concentration leads to enhancer activation and expression of downstream target genes. At the same time, zero-order ultrasensitivity also implies that a strongly repressed state is relatively stable against inappropriate activation. For example, the role of Polycomb repression, found at important developmental genes, could be to keep enhancers in the repressed regime until they are activated.

In summary, a model in which poised enhancers are actively balanced between activators and repressors could provide a mechanism to explain the ultrasensitive response of enhancers to patterning signals. This could explain the widespread occurrence of a distinct poised enhancer state during tissue patterning. Since the model makes clear mechanistic predictions, it opens new avenues for further exploration and tests in the future.

## Methods

### Stock maintenance and embryo collection

The fly stock *Tl^10b^* is from Bloomington (*Tl^10b^*, Bloomington stock center, Bloomington, #30914). The *gd^7^* stock was a kind gift from Mike Levine. *gd^7^/gd^7^* females *gd^7^*/Y males and were obtained from the *gd^7^/winscy, hs-hid* stock by heat shocking 1 day-old larvae for 1 h at 37°C, followed by a second heat shock 24 h later. *T(1;3)OR60/ Tl^10b^, e^1^* females and *Tl^10b^ /TM3, e^1^, Sb^1^, Ser^1^* males were selected from the stock consisting of genotypes *Tl^10b^ /TM3, e^1^, Sb^1^, Ser^1^* and *T(1;3)OR60/ TM3, e^1^, Sb^1^, Ser^1^*. Wild-type embryos (*Oregon-R*) at 2-4 h AED were used for GAGA, and Pc ChIP-seq. Embryos were collected on apple juice plates for 2 h at 25°C from cages and then matured at 25°C for another 2 h (2-4 h after egg deposition (AED)). Embryos were crosslinked for 15 min with 1.8% formaldehyde (final concentration in water phase).

### ChIP-seq experiments

ChIP-seq experiments were performed as described (He et al. 2011; He et al. 2015) with the following modifications. Per ChIP, ~100 mg embryos were used. After incubation of magnetic beads with antibodies, H3K27ac ChIP samples were washed 3 times by rotating tubes for 3 min at 4°C to reduce background. The antibodies for ChIP-seq were generated by Genscript (Dl aa 39-346, Mad aa 147-455, full-length Zen, Zld 211 aa near C-term, GAGA aa 163185-42), by Covance (Twi aa 340-490 or were commercially available: H3K27ac (Active motif, #39133), H3K4me1 (Active motif, #39635), H3K27me3 (Active motif, #39155), Pc (Santa Cruz, #sc-25762). *Tl^10b^* embryos were used for ChIP-seq for Dl, Twi, Zld, H3K27ac, H3K4me1, and H3K27me3; wild type embryos for GAGA and Pc; and *gd^7^* embryos for Mad, Zen, Zld, H3K27ac, H3K4me1, and H3K27me3.

### Library preparation

Different combinations of library preparation kits and barcodes were used for ChIP-seq and mRNA-seq library preparations (see Supplemental Table S3) and libraries were prepared according to manufacturer’s instructions. ChIP-seq libraries were prepared from 5-15 ng ChIP DNA or 100 ng WCE input DNA and sequenced on the GAIIX (Illumina) or the HiSeq 2500 (Illumina).

### ChIP-seq data processing

Sequenced ChIP-seq reads were aligned to UCSC *Drosophila melanogaster* reference genome dm3 using Bowtie v1.1.1 (Langmead et al. 2009), allowing up to two mismatches and retaining only uniquely aligning reads. Aligned reads were extended to the sample’s estimated fragment size using the *chipseq* Bioconductor library (Huber et al. 2015).

Replicates of genotype-specific WCE input samples for *Tl^10b^* and *gd^7^* were merged, and these merged WCEs were used for enrichment calculations and peak calling.

Transcription factor enrichments within each enhancer were calculated within a 201 bp window centered at the transcription factor’s ChIP-seq signal summit. Enrichment calculations were normalized for both differences in read count and estimated fragment size between ChIP and WCE samples. Histone modification enrichments were calculated similarly, but using a 1001 bp window centered on the enhancer region. The replicates for each transcription factor and histone modification with the highest median enrichment were used for further analysis.

### Normalization of histone modification ChIP-seq data

Fold-change in ChIP-seq enrichments of H3K27ac, H3K4me1 and H3K27me3 between *Tl^10b^* and *gd^7^* were normalized to account for differences in ChIP efficiency. The normalization factor for each histone modification was determined by the median fold-change in ChIP-seq enrichment at MACS2 peaks that were detected in both mutant embryos.

### mRNA-seq experiments

Total mRNA was extracted from 50-100 mg non-crosslinked 2-4 h AED *gd^7^* (three replicates) and *Tl^10b^* (two replicates) embryos using the Maxwell Total mRNA purification kit (Promega, Madison, WI, #AS1225) according to manufacturer’s instructions. PolyA-mRNA was isolated using DynaI oligo(dT) beads (Life Technologies, Carlsbad, CA, #61002). Libraries were prepared following the instructions of the TruSeq DNA Sample Preparation Kit (Illumina, #FC-121-2001) and sequenced on the HiSeq 2500 (Illumina).

### mRNA-seq data processing

mRNA-seq reads were aligned against the FlyBase r5.57 genome and gene annotations using Tophat2 v2.0.14 (Kim et al. 2013). Cufflinks v2.2.1 (Hemavathy et al. 1997) was used to obtain transcript abundance (FPKM).

### Data analysis

#### List of known DV enhancers

A list of known DV enhancers was assembled from the literature. Enhancers were only included if *lacZ* reporter assays were shown confirming a DV-biased expression pattern. The full list of known DV enhancer regions and the respective publication that shows the staining of the enhancer’s *lacZ* reporter assay can be found in the Supplemental Material and Supplemental Table S1. A target gene was assigned to DV enhancers only if the enhancer’s expression domain overlapped and resembled the gene’s expression domain. For enhancers identified by Kvon et al. (2012) and Ozdemir et al. (2011), published mRNA *in situ* hybridization data from the BDGP database were used for this purpose. For some of these enhancers, no target gene was identified with confidence and thus those enhancers were not included in mRNA-seq analysis shown in Supplemental Fig. S1 (see Supplemental material for more information).

#### Active enhancers, uninduced enhancers and closed regions

Active enhancers are MEs in the mutant *Tl^10b^* and DEEs in *gd^7^* embryos. Uninduced enhancers are MEs in *gd^7^* embryos and DEEs in *Tl^10b^* embryos. A total of 100 “closed regions” were randomly selected from published DHS regions (Thomas et al. 2011), which were only active at stage 14 and not in any of the earlier stages. The “closed regions” were also required to overlap with peaks from published H3K27ac 14-16 AED h in wild type embryos (modENCODE ID:4120) (Contrino et al. 2012). DHS regions that overlapped with a TSS (2 kb centered on a TSS) were excluded from the selection.

#### ChIP-seq binding profiles at single genes

Single gene profiles of histone modifications show ChIP-seq enrichment values over input calculated using a 501 bp sliding window. Transcription factor profiles are shown in reads per million.

#### Distance to putative PREs

Putative PREs were defined as regions that result from overlapping Pc and GAGA peaks (min 50 bp overlap) from ChIP-seq in wild-type 2-4 h AED embryos. Overlapping regions were combined to one putative PRE region. For Zld-bound regions, peaks were called by MACS2 on the wild-type Zld ChIP-seq sample and filtered for those with Zld binding of at least 2-fold over background in either *gd7* or *Tl^10b^*. Enrichment of H3K27me3 was calculated for each Zld peak in a region 1,000 bp centered at the peak summit. Both known enhancers and Zld regions were divided into H3K27me3 “low” and “high” groups based on an enrichment threshold of two-fold below or above input, respectively. Coordinates for putative PREs can be found in Supplemental Table S2 and distances of known DV enhancers to the closest putative PRE can be found in Supplemental Table S1.

#### DNase I hypersensitivity at known DV enhancers

Average DHS signal per base (Thomas et al. 2011) was calculated for all known enhancers at each of the five embryonic stages by summing the number of DHS reads that overlap each enhancer and dividing by the enhancer’s width in base pairs.

## Data access

ChIP-seq and mRNA-seq data are available from the NCBI Gene Expression Omnibus under the accession number GSE68983 http://www.ncbi.nlm.nih.gov/geo/query/acc.cgi?token=kdenmekojhybrkl&acc=GSE68983. In addition, a Linux virtual machine containing all raw data, processed data, analysis software and analysis code is available via Amazon Web Services. See http://research.stowers.org/zeitlingerlab/data for details.

## Acknowledgements

We thank Robb Krumlauf, Chris Rushlow, Robin Fropf, Malini Natarajan and Arnob Dutta for feedback on the manuscript, Kai Chen for providing the Pc ChIP-seq data, Cindi Staber for providing *Tl^10b^* mRNA-seq and Sam Meier for initial help with data analysis. We thank Mark Miller for scientific illustration. This work was funded by the NIH New Innovator Award 1DP2 OD004561-01 to J.Z. and the Stowers Institute for Medical Research. N.K.’s contribution was part of her PhD thesis with the Open University, UK.

